# Long-duration and non-invasive photoacoustic imaging of multiple anatomical structures in a live mouse using a single contrast agent

**DOI:** 10.1101/2022.05.17.491718

**Authors:** Anjul Khadria, Chad D. Paavola, Yang Zhang, Samuel P. X. Davis, Patrick Grealish, Konstantin Maslov, Junhui Shi, John M. Beals, Sunday S Oladipupo, Lihong V. Wang

## Abstract

Long-duration *in vivo* simultaneous imaging of multiple anatomical structures is useful for understanding physiological aspects of diseases, informative for molecular optimization in preclinical models, and has potential applications in surgical settings to improve clinical outcomes. Previous studies involving simultaneous imaging of multiple anatomical structures, e.g., blood and lymphatic vessels as well as peripheral nerves and sebaceous glands, have used genetically engineered mice, which require expensive and time-consuming methods. Here, an IgG4 isotype control antibody is labeled with a near-infrared dye and injected into a mouse ear to enable simultaneous visualization of blood and lymphatic vessels, peripheral nerves, and sebaceous glands for up to 3 hours using photoacoustic microscopy. For multiple anatomical structure imaging, peripheral nerves and sebaceous glands are imaged inside the injected dye-labeled antibody mass while the lymphatic vessels are visualized outside the mass. The efficacy of the contrast agent to label and localize deep medial lymphatic vessels and lymph nodes using photoacoustic computed tomography is demonstrated. The capability of a single injectable contrast agent to image multiple structures for several hours will potentially improve preclinical therapeutic optimization, shorten discovery timelines, and enable clinical treatments.

## Main

Concurrent long-duration live animal imaging of different anatomical structures, including blood and lymphatic vessels, peripheral nerves, and sebaceous glands can underpin an improved understanding of disease progression, study effects of therapeutics, and guide molecular optimization in preclinical models.^[1–3]^ Fluorescence microscopy has been shown to simultaneously image the above-mentioned anatomical structures using mice genetically engineered with fluorescent proteins. However, such methods are expensive, time-consuming, and are not currently translatable clinically.^[4–6]^ The discovery of lymphatic vessels in the mouse and human brain has made the need for long-duration simultaneous imaging of lymphatic vessels and other anatomical structures at cellular-level resolution ever more critical to better understand the brain-related diseases and optimize new therapies.^[7,8]^ Previous work involving preclinical imaging of lymphatic vessels has used standalone dyes such as Evans blue or indocyanine green (ICG), which are not photostable, get absorbed by the blood vessels, and cannot be used for long-duration imaging.^[3,9,10]^ The clearance of these dyes in less than 30 – 60 minutes after injection is well documented in both preclinical and clinical studies.^[10,11]^ Deep lymphatic vessels in mice have been previously imaged using near-infrared fluorescence imaging; however, those studies did not image the blood vessels simultaneously and utilized Evans blue or ICG.^[12,13]^ Previous photoacoustic imaging-based studies involving simultaneous imaging of blood and lymphatic vessels have used Evans blue or ICG, or required up to five wavelengths, which is expensive and complex.^[3,10,14,15]^ Peripheral nerves are an integral part of the nervous system linking the brain to the rest of the body. Imaging of peripheral nerves is of utmost importance, for example, to study the nervous system to develop strategies to prevent accidental injuries during surgeries.^[16,17]^ Previous work on photoacoustic imaging of peripheral nerves has been limited to only *ex vivo* studies.^[18,19]^ Studies reporting *in vitro* and *ex vivo* multi-feature photoacoustic imaging have pushed the limits of bioimaging; however, no *in vivo* studies have been reported that demonstrate simultaneous photoacoustic imaging of lymphatic vessels, peripheral nerves, and sebaceous glands along with blood vessels.^[20–22]^ Contrast agents that can facilitate long-duration and non-invasive *in vivo* photoacoustic imaging of multiple anatomical structures can benefit several preclinical and potentially clinical physiological studies and diagnoses such as peripheral neuropathy, lymphoma, vasculitis, sebaceoma, etc.^[23–25]^

Several photoacoustic-based contrast agents have been reported in recent years; however, most agents are based on tumor imaging using organic or inorganic small molecules or nanoparticles.^[26–28]^ Notably, little or no advancement has taken place in contrast agents for long-duration photoacoustic imaging of lymphatic vessels and other anatomical structures such as peripheral nerves and sebaceous glands.

In this report, we use optical-resolution photoacoustic microscopy (OR-PAM) (Figure 1a)^[29]^ to simultaneously image blood and lymphatic vessels along with peripheral nerves and sebaceous glands at cellular-level resolution in the mouse ear skin. We used photoacoustic imaging to visualize blood (label-free); however, to image lymphatic vessels, sebaceous glands, and axonal peripheral nerves, we subcutaneously injected the near-infrared light absorbing sulfo-cy7.5 dye-labeled monoclonal human IgG4 isotype control antibody. In addition, we injected the dye-labeled antibody in the mouse hind-paw to observe its deep medial lymphatic vessel and lymph node through a hand-held photoacoustic computed tomography (PACT) probe (Figure 1b).^[30]^ Dye-labeled antibodies are used widely to fluorescently label and image anatomical structures through epitope bindings; however, in this report, we use an IgG4 isotype control antibody, which does not have any specific binding epitope.^[31,32]^

**Figure 1:**
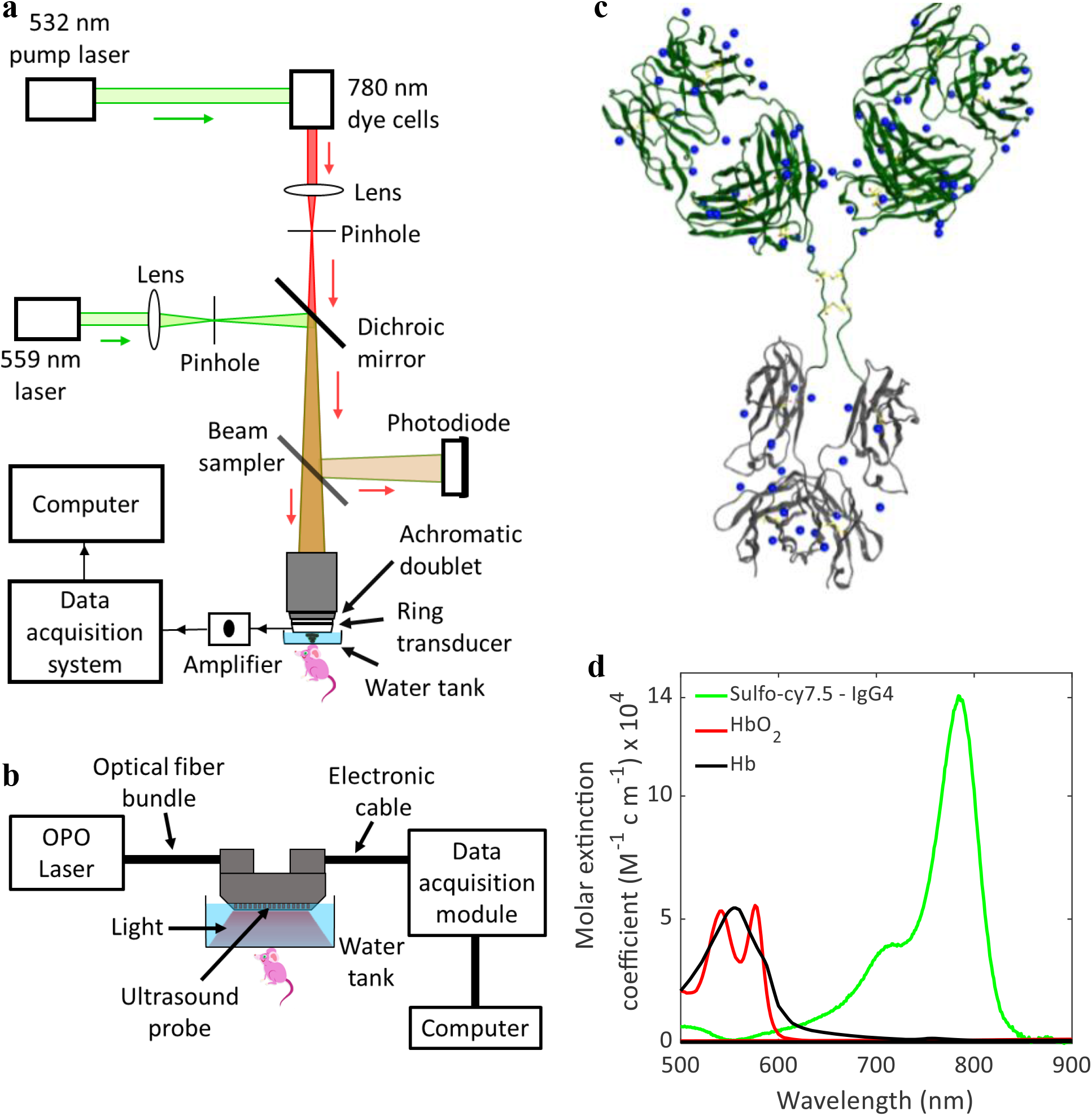
(a) Schematic of the OR-PAM system. (b) Schematic of the linear array-based PACT system. (c) Scheme of IgG4 isotype control antibody with the available sites (as blue dots) for the conjugation of sulfo-cy7.5 dye. (d) Light absorption spectra of sulfo-cy7.5 dye-labeled IgG4 antibody, oxygenated hemoglobin (HbO_2_) and deoxygenated hemoglobin (Hb).

## Results and Discussion

### Dye-labeling of the IgG4 isotype control antibody

The IgG4 antibody was labeled with the sulfo-cy7.5 dye (Figure 1c) and characterized by SEC-HPLC and MALDI mass spectrometry (see Methods). The labeled antibody was found to have approximately 4.5 dyes per molecule and exhibited size exclusion behavior consistent with a monomeric, well-behaved antibody (Figure S1 in SI). The antibody has no specific antigen binding and incorporates S228P/L234A/L235A sequence changes in the Fc region to reduce immune effector function.^[33]^ While the current study does not employ any specific paratope, antibodies with specific binding could alternatively be employed to image other anatomical structures offering tissue specificity advantage as the case may be. We further characterized the dye-labeled IgG4 antibody with UV-Vis spectroscopy and found that it has an absorption maximum at 780 nm with a molar extinction coefficient of around 14,000 M^-1^cm^-1^, which is orders of magnitude higher than that of both oxygenated and deoxygenated blood at the same wavelength (Figure 1d). A higher extinction coefficient at 780 nm will ensure high contrast signals from the dye-labeled IgG4 antibody.

### Long-duration OR-PAM of lymphatic vessels in mouse ear

We performed the OR-PAM at 559 nm and 780 nm to visualize the blood and the dye-labeled antibody, respectively, based on their absorption spectra (Figure 1d). We injected the dye-labeled antibody (0.2 μL, 20 mg/mL) into the mouse ear under anesthesia and performed photoacoustic imaging using OR-PAM. We rely on the high molecular weight (∼ 147 kDa) of the antibody to drive lymphatic absorption at the point of injection since molecules greater than 20 – 25 kDa weight are predominantly absorbed by lymphatic vessels.^[34,35]^ Antibodies, due to their large size, have a slow absorption rate from the subcutaneous injection site, which we utilize to perform long-duration imaging.^[36,37]^ The lymphatic vessels continuously absorbed the dye-labeled antibody over the first 3 hours of imaging, leading to the visualization of new lymphatic vessels while the mouse was under anesthesia (Figure 2a, Movies V1 and V2). After the first 3 hours of imaging, the mouse was allowed to recover from anesthesia and kept in an enclosure until the injection site was reimaged 6 hours’ post-injection. At 6 hours, we did not observe any lymphatic vessels stained with the dye-labeled antibody due to enhanced lymphatic clearance from the increased muscle-mediated movement of the ear in the conscious state.^[38]^ To verify that enhanced absorption of the antibody occurs while the mouse was moving, we reduced the initial anesthesia time to 60 minutes and revived the mouse for only 30 minutes, then imaged it again under anesthesia (Figure 2b). Under these experimental conditions, we did not observe any lymphatic vessels around the injection site at 90 minutes.

**Figure 2:**
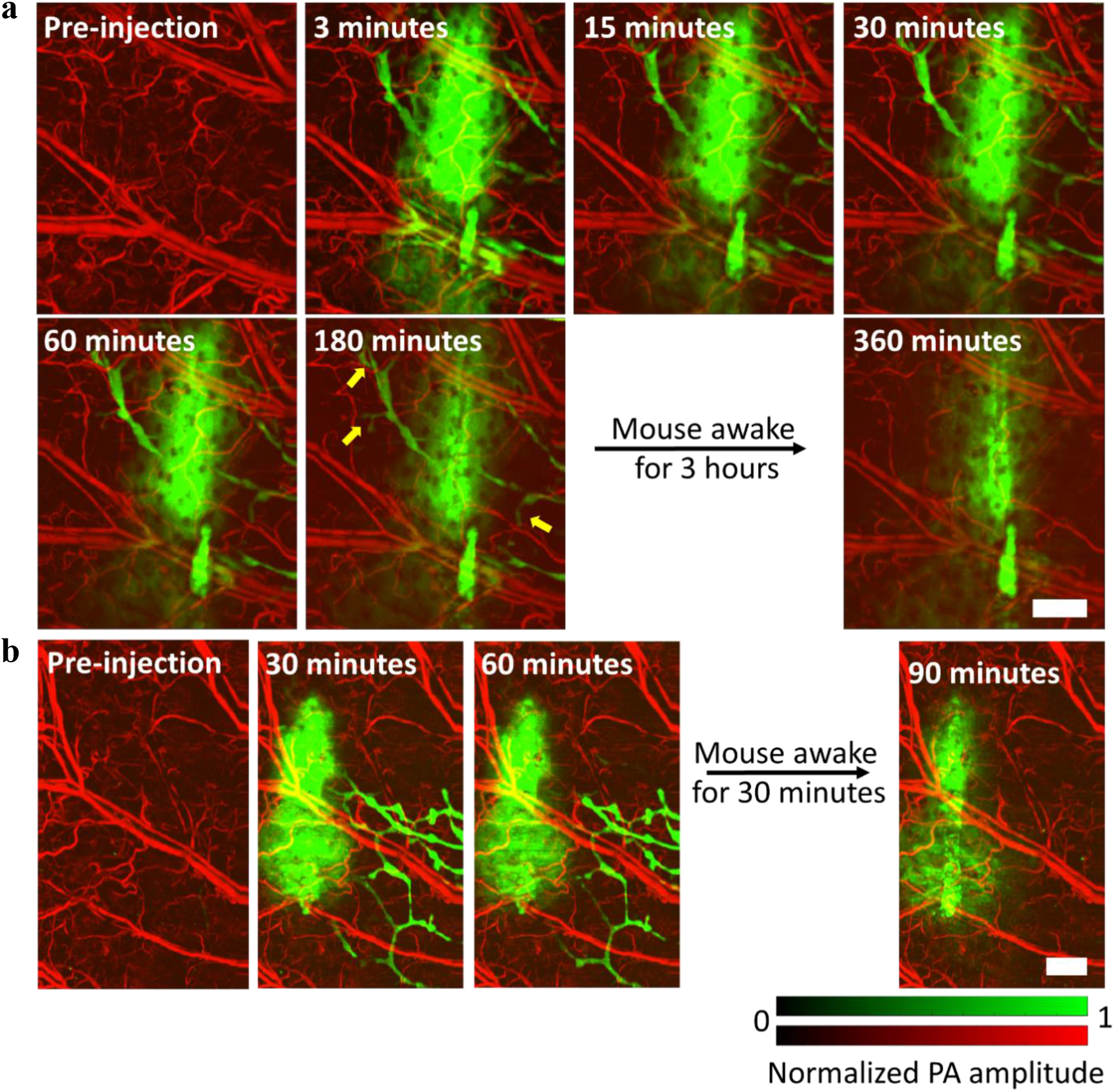
OR-PAM of lymphatic vessels in mouse ear. (a) Visualization of lymphatic vessels for up to 180 minutes upon injection of sulfo-cy7.5 dye labeled IgG4 antibody. (b) Imaging of the dye-labeled IgG4 antibody performed in the mouse ear while keeping the mouse under anesthesia for a shorter period. Yellow arrows show lymphatic vessels lighting up. PA, photoacoustic. Scale bars, 500 μm.

### Multiple anatomical structures visualizations in mouse ear

Our approach to visualize multiple anatomical features is based on imaging the peripheral nerves and sebaceous glands inside the injected dye-labeled antibody mass while visualizing the lymphatic vessels outside the mass. In our system, the 780 nm light that we used to detect the sulfo-cy7.5 dye-labeled IgG4 antibody has a depth of focus of about 300 μm, which covers the thickness of an average mouse ear (200 μm – 250 μm). The maximum amplitude projection (MAP) of the complete three-dimensional (3D) photoacoustic image shows only the injected mass of the dye-labeled antibody and the lymphatic vessels outside the mass (Figure 3a). The sebaceous glands and peripheral nerves are not clearly visible due to excess background signals (from the dye-labeled antibody) in the MAP image. We inspected two-dimensional (2D) sections at various depths to avoid the excess dye-labeled antibody background signals and distinctly visualize the axonal peripheral nerves and sebaceous glands (Figure 3b). We further performed image segmentation (see Methods) to digitally label the different structures (Figure 3c). Peripheral nerves and arteries are aligned together in mouse skin, a feature that enabled us to identify the nerves in our images.^[6,39]^ The circular structured sebaceous glands were noticeably visible in the images. The visualization of sebaceous glands in the mouse ear could be helpful in several types of skin studies.^[40]^ Many biological phenomena or disorders related to sebaceous glands, such as sebaceous adenoma, sebaceoma, sebaceous gland hyperplasia, sebaceous carcinoma, folliculosebaceous cystic hamartoma, etc. can be studied by imaging the size and shape of sebaceous glands as well as their effects on surrounding vessels.^[41]^ The major blood vessels in the mouse ear are also seen in Figure 3b, suggesting that the dye-labeled IgG4 antibody encapsulated the vessels, creating a contrast as the antibody is too large to be absorbed through the pores of the blood microvasculature.

**Figure 3:**
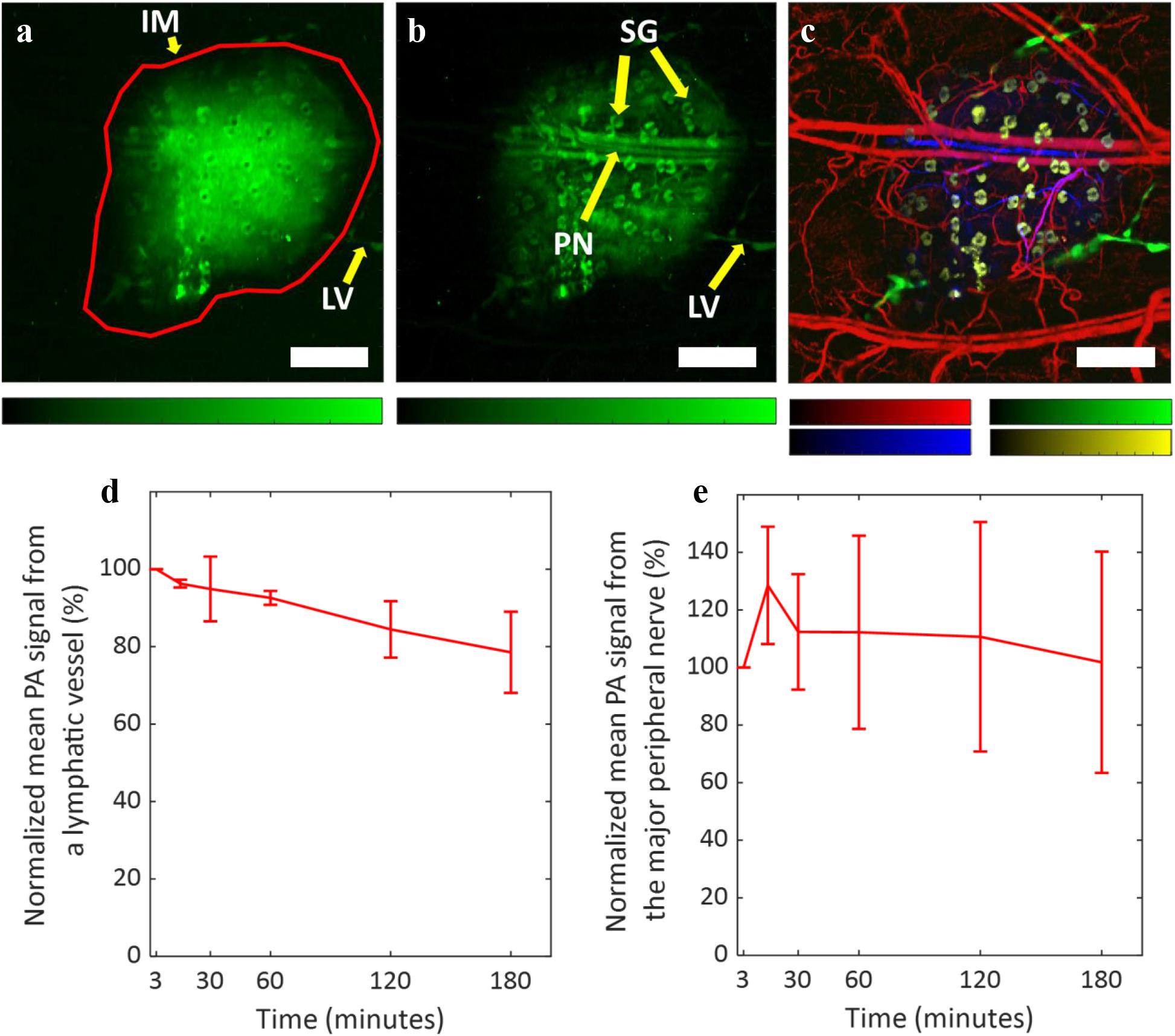
OR-PAM of multiple anatomical structures. (a) MAP image of the complete dye-labeled antibody injected in mouse ear depicting the injected dye-labeled antibody mass and the lymphatic vessels outside the mass. (b) MAP image of the dye-labeled antibody in mouse ear after inspection of 2D sections in the 3D volume to remove background signals. The sulfo-cy7.5 dye-labeled IgG4 antibody stains axonal peripheral nerves, sebaceous glands, and the major blood vessels inside the injected dye-labeled antibody mass, and the lymphatic vessels outside the mass. (c) Different anatomical structures labeled in pseudo colors through image segmentation. Blood vessels are colored as red (559 nm), lymphatic vessels as green (780 nm), sebaceous glands as yellow (780 nm), and axonal peripheral nerves as blue (780 nm). (d) The mean photoacoustic signal from a single lymphatic vessel. (e) The mean photoacoustic signal from the major nerve. Data represent mean ± standard deviation (n = 3). IM, injected mass; LV, lymphatic vessel; PN, peripheral nerve; SG, sebaceous gland. All color bars represent normalized photoacoustic amplitude and range from 0 to 1. Scale bars, 500 μm.

Upon quantitative comparison of the mean photoacoustic signal from a single lymphatic vessel that remains visible throughout the imaging time, we found that the photoacoustic signal does not fade away significantly (∼ 20% decrease) even 3 hours after the injection under anesthetized conditions (Figure 3d). As mentioned above, the widely used lymphatic contrast agents such as ICG or Evans blue get cleared away in less than 1 hour.^[10,11]^ These results validate the efficacy of the large-sized dye-labeled monoclonal antibody for long-duration lymphatic imaging. The peripheral nerves were also visible for up to 3 hours (Figures 3e and S2 in SI) without a significant decrease in the mean photoacoustic signal. Note that some signals were detected even after 50 days of injection (Figure S3 in SI).

### Long-duration deep lymphatic vessel visualization

We injected the dye-labeled antibody in the mouse hind paw and performed imaging away from the site of injection through PACT at 780 nm and 920 nm in the leg and thigh areas where the deep medial lymphatic vessel and lymph node are located to observe them (Figures 4a, 4b, and S4 in SI, Movie V3).^[12]^ Although the pre-injection and post-injection images at 780 nm were sufficient to establish the dye-labeled antibody uptake by the lymphatic vessel and lymph node, we performed imaging at 920 nm to confirm that no blood leakage occurred during needle insertion while performing the injection. We chose 920 nm to perform imaging of blood because the extinction coefficient of hemoglobin in blood in the NIR region (without overlapping with the absorption spectrum of the dye-labeled IgG4 antibody) is highest at around 920 nm, which ensures more photoacoustic signal from the blood.^[42]^ The images in Figure 4 were processed by vessel segmentation (see Methods). We performed imaging for up to 3 hours following injection to continuously observe bright signals from the lymphatic vessels at a depth of 2 – 4 mm from the surface of the mouse skin, thus proving the long-duration efficacy of the method (Figures 4b and S4 in SI). The dye-labeled antibody does not absorb any light at 920 nm, but the blood absorbs significant light. We did not perform any multi-feature imaging using PACT in deep tissues because of two reasons. (1) The sebaceous glands are present only in the skin for which high resolution-based OR-PAM is sufficient. (2) We have no pre-information about the nerves in deep tissues to correctly identify them such as we had for mouse skin where the peripheral nerves and arteries are aligned together.^[6,39]^

**Figure 4:**
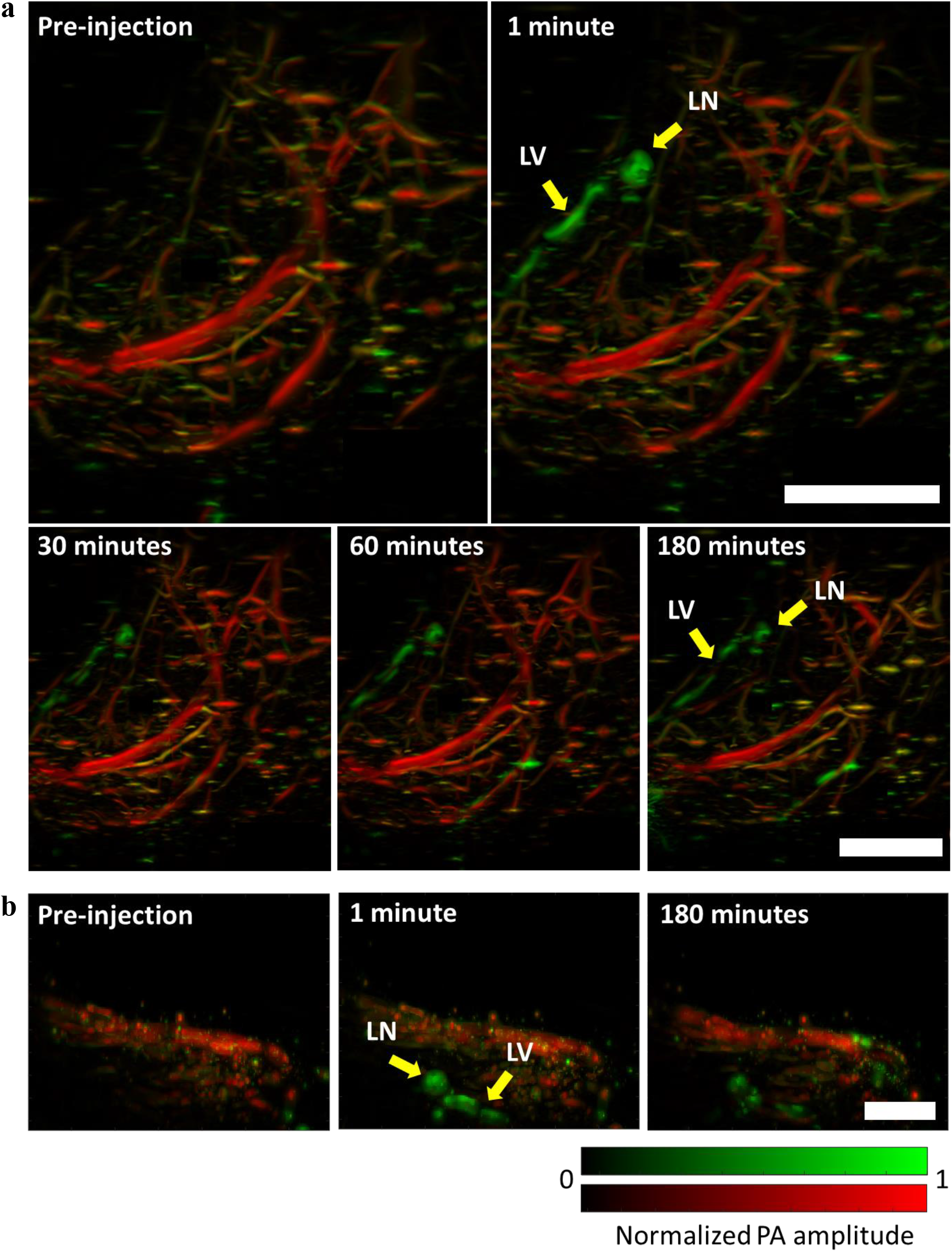
Long-duration photoacoustic imaging of deep lymphatic vessels in mouse. (a) Top-view images of deep medial lymphatic vessel and lymph node (green), and blood (red) in mouse leg and thigh taken through PACT. (b) Side-view images of deep medial lymphatic vessel (green) and blood (red) in mouse leg and thigh taken through PACT. LV, lymphatic vessel; LN, lymph node; PA, photoacoustic. Scale bars, 0.4 cm.

These results show that the dye-labeled IgG4 antibody can be used for long-duration photoacoustic imaging of superficial and deep lymphatic vessels. Long-duration imaging of deep lymph nodes and lymphatic vessels could be a powerful tool for visualizing deep tumors during cancer metastasis.

## Conclusions

In this study, we used IgG4 monoclonal antibody labeled with near-infrared sulfo-cy7.5 dye to perform photoacoustic imaging of multiple anatomical structures in mouse ear skin and deep medial lymphatic vessels in the leg and thigh. The slow clearance rate of the large-sized antibody allows long-duration photoacoustic imaging of the anatomical structures, which is otherwise difficult to perform with the range of existing contrast agents. The long-duration multi-anatomical structure labeling feature of the dye-labeled antibody can be used in several types of preclinical studies involving studies of diseases and drug therapeutics. It also has the potential to be clinically translatable. The antibodies can be further designed to bind to specific antigens to enable long-duration imaging of deep and shallow individual anatomical structures *in vivo* at high resolution, which is otherwise difficult to image due to a lack of sufficient injectable contrast agents, especially for photoacoustic imaging.

## Methods

### Sulfo-cy7.5 dye labeling of human IgG4 isotype control antibody

An IgG4 isotype antibody was labeled with Sulfo-Cyanine-7.5 dye using NHS ester chemistry (66320, Lumiprobe). The antibody was prepared at approximately 10 mg/mL in 90 mM carbonate, 9 mM phosphate buffer with 125 mM sodium chloride at pH 8.3. The dye was dissolved in a one-tenth volume of carbonate buffer immediately before adding to the antibody at a 10 equivalent excess and incubated at 25°C for 4 hours. The labeled antibody was isolated from the excess dye on a size exclusion chromatography column (Superdex S200, Cytiva) with a mobile phase of 1x PBS, pH 7.2 at 1 mL/min. The degree of labeling was determined using MALDI-MS to be ∼ 4.5 dyes per antibody. The antibody was concentrated to ∼ 30 mg/mL using a 15 mL spin concentrator (Millipore) with a 100 kDa MWCO membrane. Samples of labeled and unlabeled antibody (5 – 10 μg) were run on a HPLC (Agilent 1260 Infinity II) using an Agilent AdvanceBIO SEC 300Å 2.7 mm column (PL1580-3301, Agilent) at 1 mL/min in a mobile phase of 1x PBS; pH 7.2 (20012-027, GIBCO) with peaks detected by absorbance at 214 nm. Total run time was 7 minutes.

### Measurement of the extinction coefficient of sulfo-cy7.5 dye-labeled IgG4 antibody

The absorbance of the sulfo-cy7.5 dye-labeled IgG4 antibody (1 μM antibody, 4.5 dyes labeled per antibody) was measured using a UV-Vis spectrophotometer (Cary® 50 Bio, Varian) at 20 °C. The extinction coefficient was calculated using the Beer-Lambert law, A = εCL where A is the absorbance, ε is the molar extinction coefficient, C is the dye concentration, and L is the light path length (1 cm). The molar extinction coefficient values for oxygenated and deoxygenated blood were taken from the compilation of Scott Prahl.^[42]^

### Photoacoustic microscopy (PAM) system design

We employed a spherically focused ring-shaped transducer (central frequency = 42 MHz, f-number = 1.67, Capistrano Labs) in the PAM, which is equipped with two lasers of 559 nm and 780 nm optical wavelengths. Light pulses of both wavelengths were used to irradiate the same point successively with microseconds delay. The light beam from a 559 nm Nd:YAG laser (BX2II, Edgewave) was focused on a 254 μm diameter orifice (3928T991, McMaster-Carr) for spatial filtering. A dye laser (Credo, Sirah) pumped by a 532 nm Nd:YAG laser (IS80-2-L, Edgewave GmbH) was employed to generate the 780 nm light beam. Styryl 11 dye (07980, Exciton) in 200 proof ethanol (MFCD00003568, Koptec) was circulated in the dye laser at 18 °C. The 780 nm light beam was combined with the 559 nm light beam through a dichroic mirror (M254C45, Thorlabs). Before the combination, the 780 nm beam was focused on a second pinhole (3928T991, McMaster-Carr) for spatial filtering. A separate pinhole for the 780 nm light beam was used to increase its depth of focus (∼ 300 μm) with a lower resolution and also to correct for the focal length difference caused by light dispersion in the immersion liquid. An achromatic doublet (AC080-020-A, Thorlabs) was used to focus the combined beam on the sample. Some amount of light was collected by a photodiode (PDA36A, Thorlabs) with the aid of a beam sampler to correct for laser fluctuations. We raster scanned the animal using a stepper motor-based two-dimensional scanner (PLS-85, Physik Instrumente) controlled by a customized LabVIEW program with an FPGA (PCIe-7841, National Instruments). The data was acquired using a digitizer (ATS 9350, AlazarTech) at 500 MS/s.

### Nano-liter injection for photoacoustic imaging

We used 31-gauge needles (7803-03, Hamilton) fitted in microliter syringes to inject antibody solutions at volumes of 0.10 μL and 0.2 μL in the mouse ear. To perform the rapid and controlled injection, the syringes were fitted onto a syringe dispenser (PB600, Hamilton). A 5 μL (7634-01, Hamilton) syringe was used to perform the injections.

### Photoacoustic computed tomography (PACT) system design

The PACT system consists of a laser source for optical illumination, an ultrasound linear array for recording the photoacoustic signals, a motor to perform a linear scan, a data acquisition (DAQ) module to digitize the signals, and a processing system for reconstructing the image (Figure 1b). We used an optical parametric oscillator (OPO) laser (SplitLight EVO III - 100, Innolas) to deliver light at wavelengths of 780 nm and 920 nm to the target through an optical fiber bundle. The linear probe with a bandwidth of 13 – 24 MHz (MS 250, VisualSonics) was mounted on the stepper motor (PLS-85, Physik Instrumente) and connected to a Verasonics Vantage 256 system (Verasonics). It featured a 14-bit analog-to-digital converter (ADC) dynamic range and sampling frequencies up to 62 MHz. The PACT system operated at 100 Hz in trigger mode (triggered by a laser) to acquire 600 frames in 6 seconds. We used the universal back-projection algorithm to reconstruct the images.^[43]^

### Animal experiments

We performed all imaging experiments on mice using protocols approved by IACUC at the California Institute of Technology. We used Hsd:Athymic Nude-Fox1^nu^ mice aged 4 to 12 weeks (Envigo) in all experiments while maintaining their body temperatures at 37 °C during imaging. All the mice were imaged under isoflurane anesthesia (1.25 – 1.50 % isoflurane in the air at a flow rate of 1 L/min).

#### OR-PAM

The mouse ear was imaged at a 4 kHz A-line rate with a fast axis (2.5 μm step size, 1100 steps) and a slow axis (5 μm step size, 800 steps), resulting in a total scanning time of 220 seconds. To image the dye-labeled IgG4 antibody formulations in a mouse ear, we took a pre-injection image of the mouse ear before performing a sub-microliter injection of the required antibody formulation. Then images were taken at 3 minutes, 15 minutes, 30 minutes, 60 minutes, and 180 minutes’ time points, during which the mouse was maintained under anesthesia. The imaging time-point was defined as the mid-point between imaging start and finish. After 180 minutes, the mouse was awakened and kept in its cage with food and water. The mouse was anesthetized and reimaged at 6 hours’ time point post-injection.

#### PACT

The 3D images were acquired by scanning the ultrasound probe (step size = 50 μm) with a total time of 6 seconds. The backside of the mouse leg and thigh regions where the deep medial lymphatic vessel is located was imaged using light with wavelengths of 920 nm and 780 nm. Then the dye-labeled antibody (50 μL, 20 mg/mL) was injected into the hind-paw and the image was immediately acquired (within 1 minute). Subsequent images were acquired at 15 minutes, 30 minutes, 1 hour, 2 hours, and 3 hours after injection during which the mouse was maintained under continuous anesthesia at 37 °C.

### Image segmentation

To segment the different anatomical structures, we applied a series of operations to all sections where relevant features were visible. We first applied total variance denoising,^[44]^ then used Gaussian filters and morphological top-hat filters to select structures of relevant scale. The sebaceous glands were masked in the first image using a Canny edge detection algorithm^[45]^ to detect the boundaries of the glands, which were filled with a morphological closing, and cleaned with a morphological dilation. The lymph vessels were isolated by similarly masking the central absorption region and using the complimentary mask. After the initial contrast-enhancing operations, the peripheral nerves and the sebaceous glands in the second image were manually labeled.

### Quantification of mean photoacoustic signal from the lymphatic vessel and the peripheral nerve

The areas covered by the major peripheral nerve or a lymphatic vessel (that is visible throughout the time series) in the MAP images were roughly selected (after passing the images through an average filter of size 3 × 3 pixels). Then the images were thresholded by the summation of the mean and three times the standard deviation of the background amplitude to segregate photoacoustic signals from the noise. The contrast to noise ratio of resultant photoacoustic amplitudes was calculated to acquire the mean photoacoustic signal from the peripheral nerve or the lymphatic vessel. The mean of the signal at all the time points was divided by the mean at 3 minutes and then multiplied by 100 to calculate the percentage of the photoacoustic amplitude with respect to the initial time point.

### Vessel segmentation for PACT images

Post-processing of reconstructed PACT volumes was performed with MATLAB (Mathworks). Vessels were segmented using a Hessian-based multiscale vessel enhancing operator.^[46]^ The eigenvalues of the Hessian matrix (|*λ*_1_| ≤ |*λ*_2_| ≤ |*λ*_3_|) give a measure of the local curvature of the volume, which can be used to determine the probability that a pixel is part of a vessel-like structure. The Hessian at of a volume *P* at voxel ***x*** and scale *σ* is defined by equation M1,

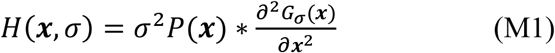

where *G*_*σ*_ is a gaussian kernel with a standard deviation *σ*. The vessel segmentation operator is then given by equation M2,

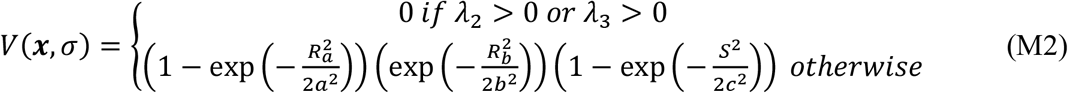

where 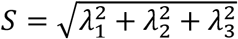 is a measure of the total curvature at ***x***, 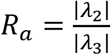 discriminates between vessel-like and plate-like structures, and 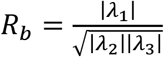 discriminates sphere-like structures. The parameters were set as *a* = 0.5, *b* = 0.5, *c* = max *λ*_3_/2. Scale invariant vessel segmentation is performed by taking the maximum of *V*(***x***, *σ*) for *σ* between 40 μm and 360 μm.

The processed 780 nm and 920 nm vessel maps were then registered. The 920 nm segmented vessels were used to mask the dye present in vasculature. Maximum projections are displayed on a logarithmic scale.

## Supporting information

Supplementary Images

## Data availability

The data that support the conclusions are mentioned in the main draft or the supplementary information.

## Author contributions

L.V.W., S.O., J.M.B, C.D.P., and A.K. conceived the project and the ideas. C.D.P. and A.K. designed the chemistry and parameters for dye labeling. P.G. labeled the antibody with the dye and characterized them. P.G. and A.K. prepared the antibody and dye buffer solutions. A.K. and K.M. designed and built the scanning photoacoustic microscope. A.K. designed and performed all the PAM experiments and analyzed the data. Y.Z. designed the PACT system and data reconstruction algorithm. A.K. and Y.Z. performed the PACT experiments. S.P.X.D. performed the image segmentation for the PAM data and vessel segmentation for the PACT data. J.S. wrote the LabVIEW software for photoacoustic data acquisition. L.V.W., S.O., and J.M.B. supervised the project. A.K. wrote the manuscript. C.D.P., Y.Z., S.P.X.D., J.M.B., S.O., and LV.W. contributed to writing the manuscript.

