## Supplementary Images for "Long-duration and non-invasive photoacoustic imaging of multiple anatomical structures in a live mouse using a single contrast agent"

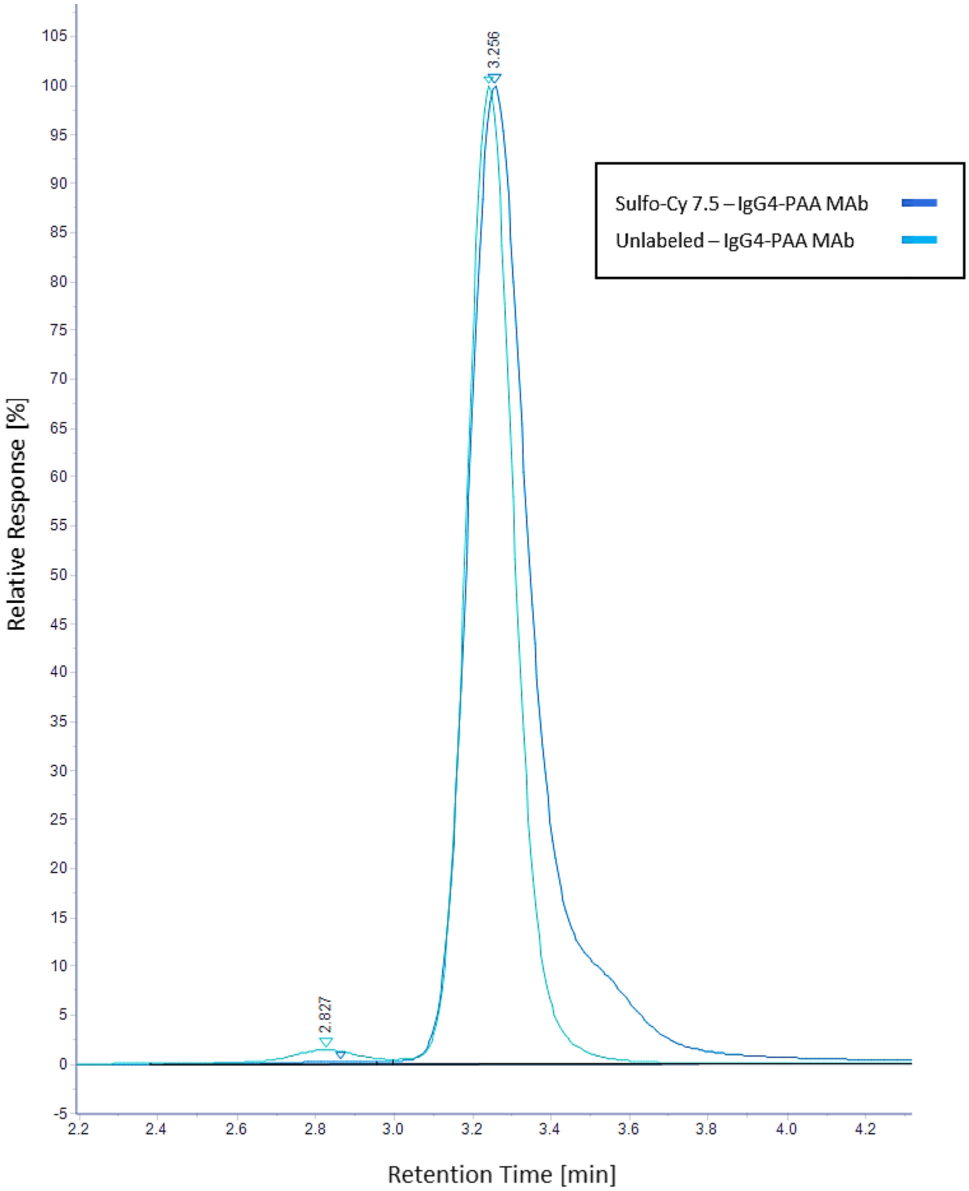


**Figure S1:** Overlay of sulfo-cy7.5 labeled (light blue) and unlabeled (dark blue) IgG4 isotype control antibody.

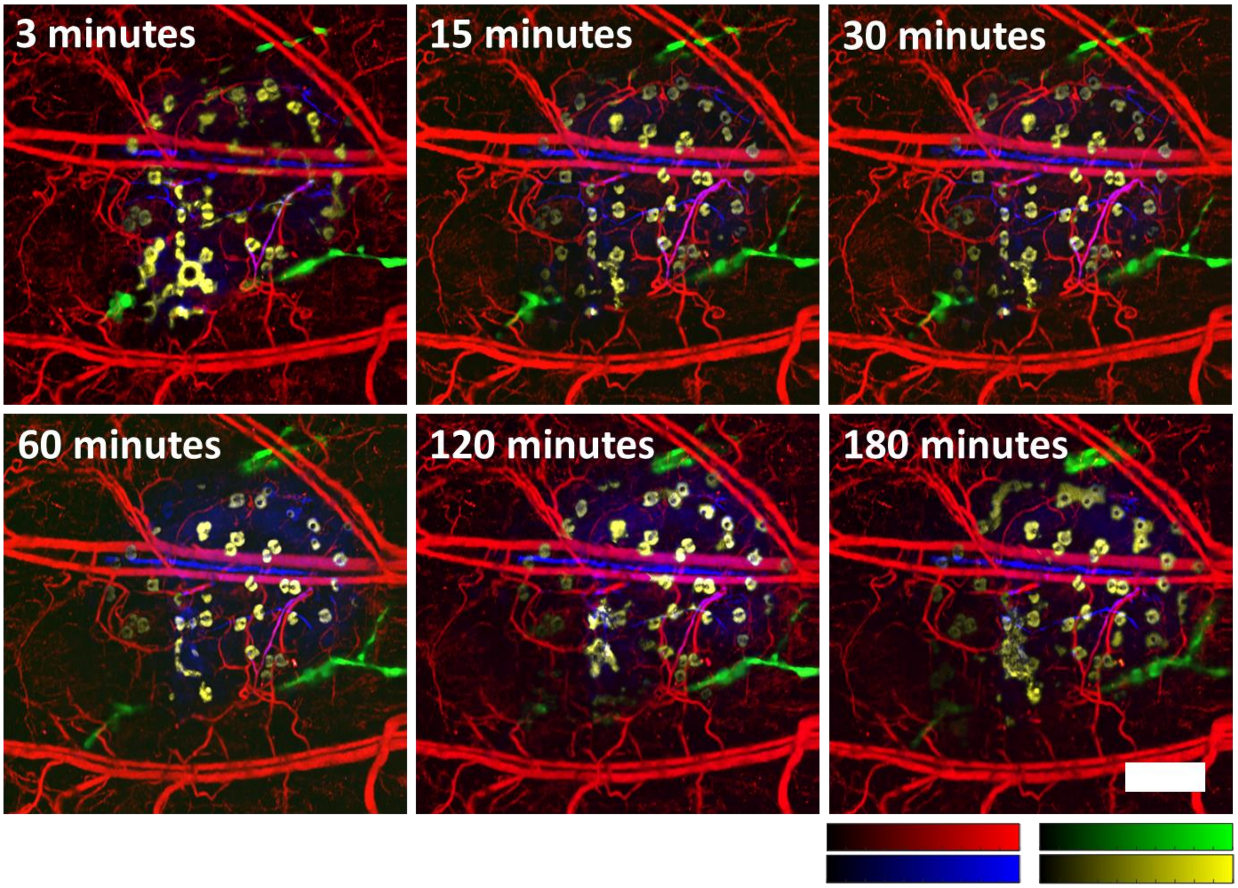


**Figure S2:** Long-duration imaging of multiple biological features in mouse ear skin after injection of dye-labeled monoclonal IgG4 isotype control antibody. All color bars represent normalized photoacoustic amplitude and range from 0 to 1. Scale bar, 500 μm.

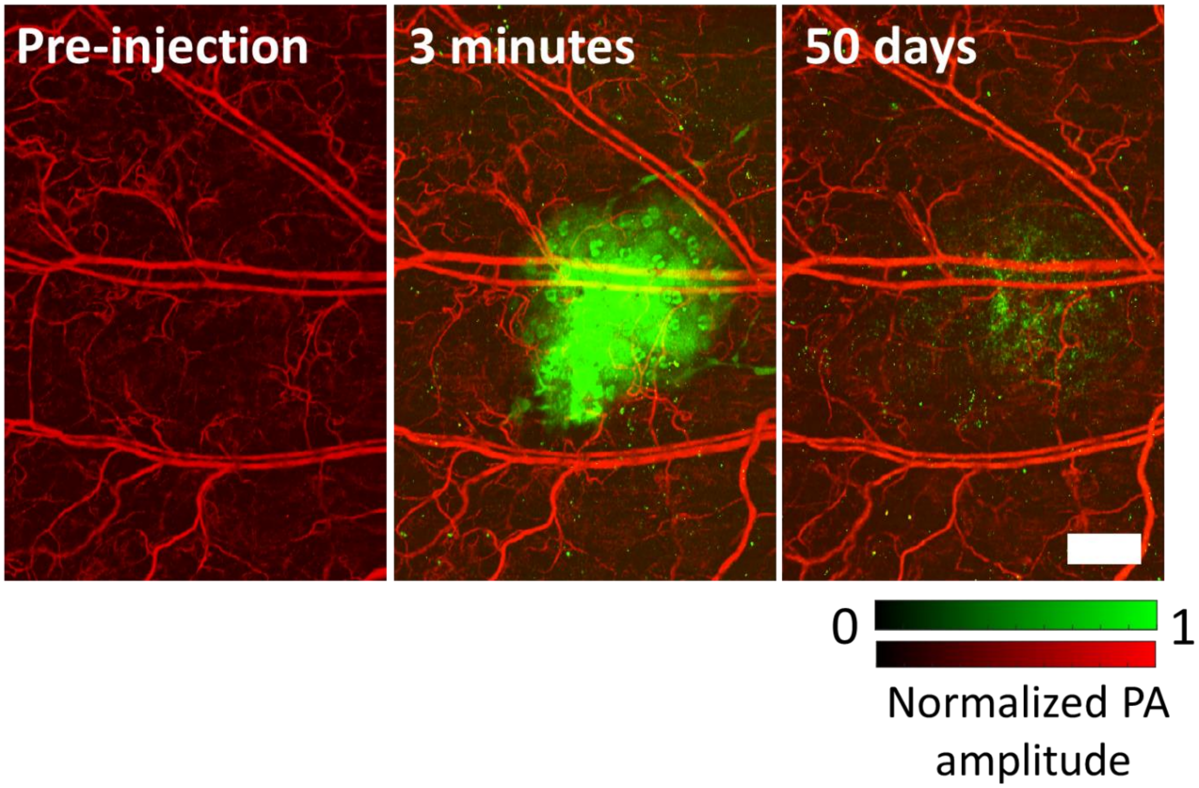


**Figure S3:** Detection of signals from dye-labeled IgG4 antibody even after 50 days of injection confirming its slow clearance rate. Scale bar, 500 μm.

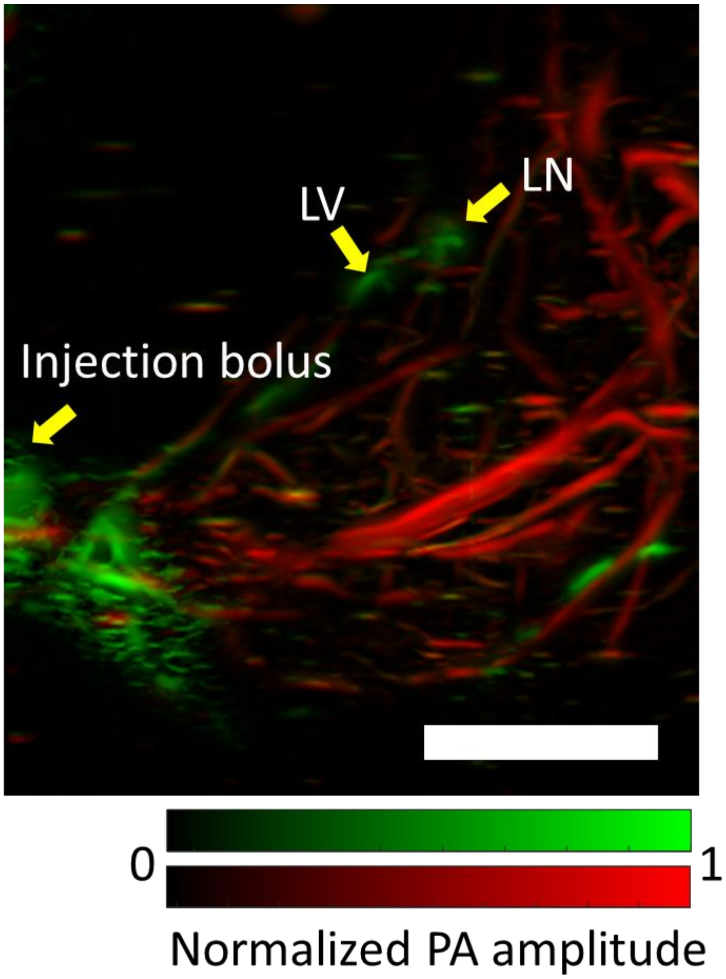


**Figure S4:** Visualization of deep medial lymphatic vessel and lymph node in the mouse leg and thigh 185 minutes after the injection of dye-labeled IgG4 antibody in mouse hind-paw using PACT. LV, lymphatic vessel; LN, lymphatic node; PA, photoacoustic. Scale bar, 0.4 cm.
